# Sensory noise increases metacognitive efficiency

**DOI:** 10.1101/189399

**Authors:** Ji Won Bang, Medha Shekhar, Dobromir Rahnev

**Author notes:** Correspondence: Dobromir Rahnev, Georgia Institute of Technology, 654 Cherry Str NW, Atlanta, GA 30332.

## Abstract

Visual metacognition is the ability to employ confidence ratings in order to predict the accuracy of ones decisions about visual stimuli. Despite years of research, it is still unclear how visual metacognitive efficiency can be manipulated. Here we show that a hierarchical model of confidence generation makes a counterintuitive prediction: Higher sensory noise should increase metacognitive efficiency. The reason is that sensory noise has a large negative influence on the decision (where it is the only corrupting influence) but a smaller negative influence on confidence (where it is one of two corrupting influences; the other one being metacognitive noise). To test this prediction, we used a perceptual learning paradigm to decrease the amount of sensory noise. In Experiment 1, seven days of training led to significant decrease in noise as well as a corresponding decrease in metacognitive efficiency. Experiment 2 showed the same effect in a brief 97-trial learning for each of two different tasks. Finally, in Experiment 3, we experimentally manipulated stimulus contrast to increase sensory noise and observed a corresponding increase in metacognitive efficiency. Our findings demonstrate the existence of a robust positive relationship between sensory noise and metacognitive efficiency. These results could not be captured by a standard model in which decision and confidence judgments are made based on the same underlying information. Thus, our study provides a novel way to directly manipulate metacognitive efficiency and suggests the existence of metacognitive noise that corrupts confidence but not the perceptual decision.

## Introduction

When faced with difficult decisions, people not only make an informed choice but can also provide a metacognitive estimate of the likelihood that their response was correct (Metcalfe & Shimamura, 1994). This judgment is usually provided in the form of a confidence rating. The ability of confidence judgments to distinguish between correct and wrong answers determines the degree of visual metacognition. High metacognitive scores suggest that confidence judgments are informative and should be trusted, while low scores suggest the opposite. Despite the importance of understanding when confidence judgments are particularly useful and when they are less so, the factors determining the quality of metacognition are still not understood.

Research into the determinants of visual metacognition has been hampered by existing measures of metacognition. Traditional metrics include the trial-to-trial Pearson correlation between confidence and accuracy (Nelson, 1984), the area under the Type 2 curve (Fleming, Weil, Nagy, Dolan, & Rees, 2010), and *type-2 d’* (Higham, Perfect, & Bruno, 2009). The quantities measured by all of these metrics increase trivially as stimulus sensitivity increases (Maniscalco & Lau, 2012). Consequently, such metrics are said to measure *metacognitive sensitivity* (Fleming & Lau, 2014): the quality of confidence ratings without regard for stimulus sensitivity.

Recently, Maniscalco and Lau (2012) developed a way to measure *metacognitive efficiency* (Fleming & Lau, 2014): the quality of confidence ratings normalized by stimulus sensitivity. Their method computes an index (of metacognitive sensitivity) *meta-d’* that can then be divided by the level of stimulus sensitivity *d’*. The resulting metric is called *M_ratio_* (Maniscalco & Lau, 2012). (Note that *meta-d’* can alternatively be normalized by subtracting *d’;* the resulting metric is called *M_diff_*.) By constructing a measure of metacognitive efficiency, the development of *M_ratio_* allows researchers to investigate metacognition independent of stimulus sensitivity.

Armed with a measure of metacognitive efficiency, we explored what factors influence metacognitive efficiency and whether it is possible to manipulate it experimentally. To do so, we turned to existing models of confidence generation. Most current models assume that confidence is based on the exact same information used to make the perceptual decision (Fetsch, Kiani, Newsome, & Shadlen, 2014; Hangya, Sanders, & Kepecs, 2016; Pouget, Drugowitsch, & Kepecs, 2016; Rahnev, Bahdo, de Lange, & Lau, 2012; Sanders, Hangya, & Kepecs, 2016). These models predict that while higher stimulus sensitivity leads to higher metacognitive sensitivity, it results in constant metacognitive efficiency. However, several newer models have included an extra level of metacognitive noise that corrupts the confidence but not the decision judgments (Berg & Ma, 2016; De Martino, Fleming, Garrett, & Dolan, 2013; Jang, Wallsten, & Huber, 2012; Mueller & Weidemann, 2008; Rahnev, Nee, Riddle, Larson, & DEsposito, 2016). We refer to these models as “hierarchical” models of confidence (Maniscalco & Lau, 2016) since they include two separate stages of noise corruption: the perceptual decision is corrupted by a first-level sensory noise, while the confidence rating is additionally corrupted by a second-level metacognitive noise (Figure 1A). Because the perceptual decision and confidence are based on different information, hierarchical models of confidence allow in principle for dissociations between metacognition and stimulus sensitivity resulting in non-constant metacognitive efficiency. Still, there has been no theoretical or empirical work on how such dissociations can be achieved.

Here we report on a counter-intuitive prediction of hierarchical models of confidence, namely that higher sensory noise should lower stimulus sensitivity but increase metacognitive efficiency. This prediction stems from the differential effect of sensory noise on stimulus and metacognitive sensitivity. Stimulus sensitivity is only corrupted by sensory noise, while metacognitive sensitivity is corrupted by *both* sensory and metacognitive noise. Therefore, increasing sensory noise is more detrimental to stimulus sensitivity than metacognitive sensitivity, resulting in higher metacognitive efficiency. Mathematically, stimulus sensitivity *d’is* the ratio of the signal and sensory noise, while *meta-d’* is the ratio of the signal and a combination of sensory and metacognitive noise. Therefore, increasing sensory noise levels has a large negative effect on *d’*but a smaller negative effect on *meta-d’*, thus leading to an increase in their ratio (that is, *M_ratio_;* Figure 1B; for a complete proof, see Methods). Importantly, a standard model based on signal detection theory (SDT), which lacks a separate metacognitive noise stage, predicts that metacognitive efficiency remains constant for different sensory noise levels (Figure 1C).

**Figure 1.**
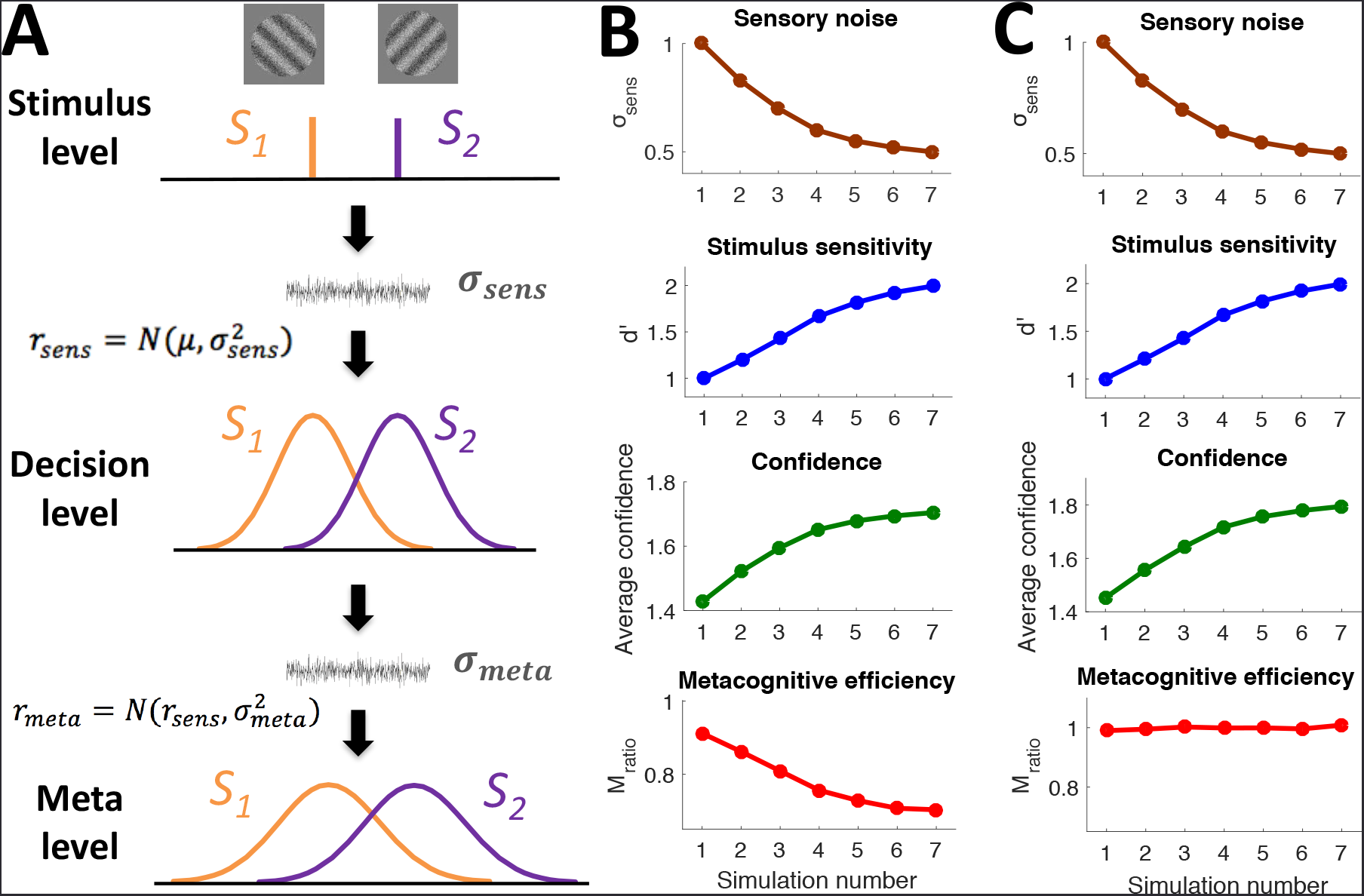
Hierarchical model of confidence. A. Generative model of confidence generation. At the stimulus level, two stimulus categories S_1_ and S_2_ (e.g., Gabor patches of counterclockwise and clockwise orientation) are presented. The stimuli are perfectly distinguishable. However, the internal representation at the decision level, r_sens_, IS corrupted by Gaussian noise σ_sens_ and thus the two stimulus categories are not perfectly distinguishable at the time of the decision. The confidence judgment is then made at the meta level based on an internal response r_meta_ that is derived from r_sens_ but is corrupted by additional noise σ_meta_. B. Depiction of the model predictions. Seven simulations with a gradually decreasing level of sensory noise, σ_sens_, show a gradual increase in sensory sensitivity d’ and confidence ratings (given on a 2-point scale such that high confidence was provided when probability of being correct exceeded 70%), but a decrease in metacognitive efficiency M_ratio_. C. Depiction of predictions made by a standard model based on signal detection theory (SDT). The SDT-based model is equivalent to the hierarchical model but lacks a metacognitive noise stage. The same decrease in sensory noise leads to similar increases in sensory sensitivity and confidence, but no change in metacognitive efficiency.

We empirically tested and confirmed the hierarchical model’s prediction that higher sensory noise leads to higher metacognitive efficiency. In two experiments, we used learning to decrease the level of sensory noise and observed a corresponding decrease in metacognitive efficiency. In a third experiment, we experimentally increased the level of sensory noise and found a corresponding increase in metacognitive efficiency. These results demonstrate that metacognitive efficiency depends on low-level stimulus characteristics and provide strong support for the existence of metacognitive noise assumed by hierarchical models of confidence.

## Results

### Experiment 1: Perceptual learning decreases metacognitive efficiency

To test the counterintuitive prediction that decreasing sensory noise leads to lower metacognitive efficiency, we employed a perceptual learning paradigm. Twelve subjects participated in a 7-day training on a visual task. Subjects performed a 2-interval forced choice (2IFC) orientation detection task in which they indicated the interval (first or second) that contained a Gabor patch (Figure 2A). Stimulus intensity was adjusted using a 2-down-1-up staircase procedure that allowed us to determine subjects intensity threshold.

Consistent with a decrease in sensory noise, training gradually decreased subjects intensity threshold (t_11_ = −5.28, *p* = .0003; one-sample t-test on the slope of change; Figure 2B). Next, we selected the same range of intensity values across all seven days of training (we used intensity values in the 35-65 percentile range; using larger percentile ranges produced similar results; see **Supplementary Results**). When considering only this range of intensity values, we observed that training increased stimulus sensitivity *d’* (t_11_ = 5.2, *p* = .0003; Figure 2B) as well as average confidence (t_11_ = 2.43, *p* = .034; Figure 2B).

**Figure 2.**
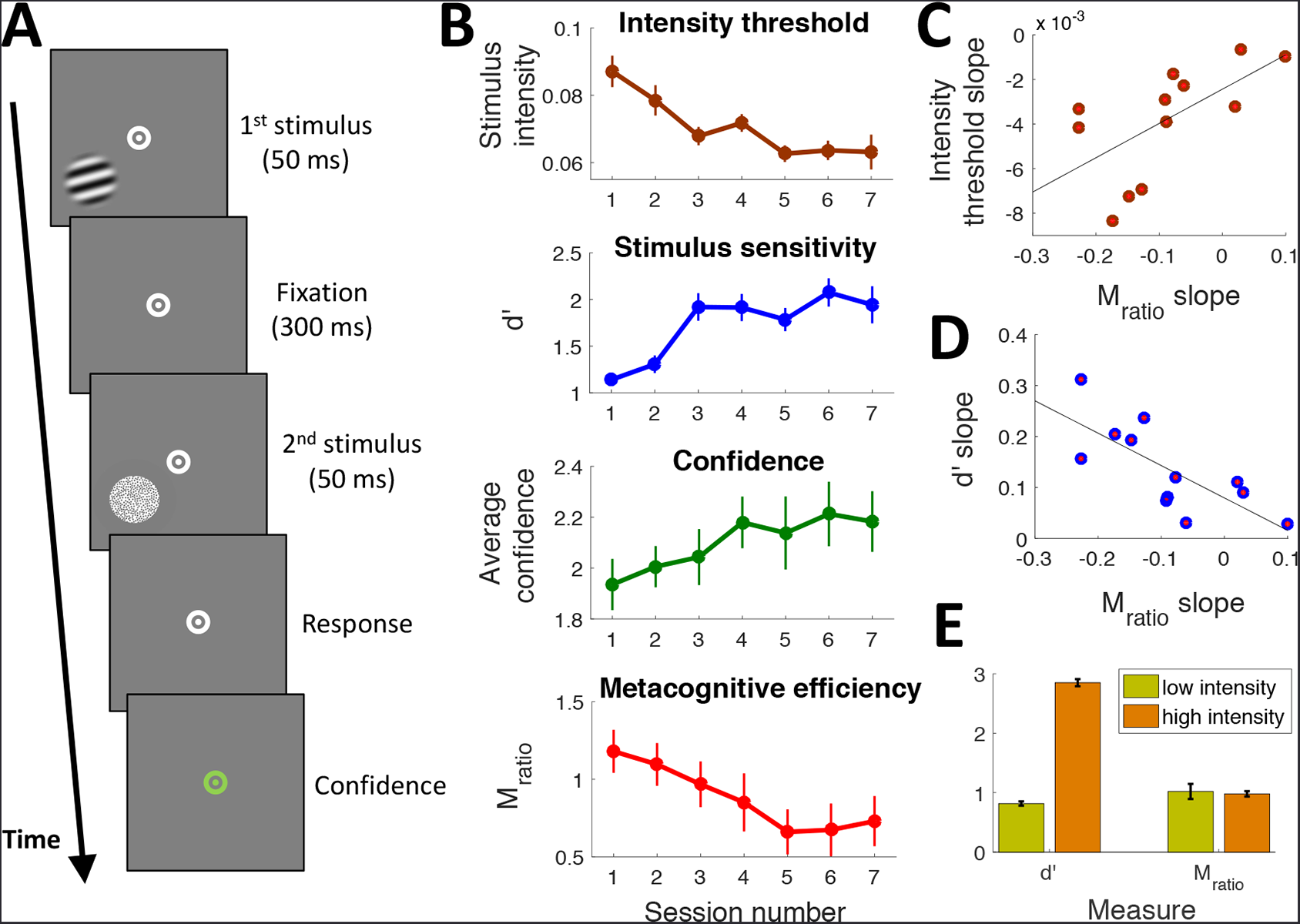
Visual training decreases metacognitive efficiency. A. Subjects performed a 2-intervalforced choice orientation detection task. Two stimuli - a target consisting of a noisy Gabor patch and a non-target consisting of pure noise - were presented in a temporal sequence. Subjects indicated the interval in which the target appeared and provided a confidence rating on a 4-point scale. B. Results of the seven days of training indicate that intensity threshold gradually decreased, while stimulus sensitivity and confidence ratings increased. Critically, as predicted by our model (Figure 1B), training decreased metacognitive efficiency. C-D. The strength of the M_ratio_ decrease on a subject-by-subject basis depended on the decrease in intensity threshold (C) and increase in stimulus sensitivity (D). E. Increased stimulus sensitivity does not automatically result in a M_ratio_ decrease. Comparing low and high intensity stimuli shows a large difference in stimulus sensitivity d’ but no difference in M_ratio_. Error bars indicate S.E.M.

Critically, as predicted by our hierarchical model of confidence, the decreased sensory noise also resulted in decreased metacognitive efficiency *M_ratio_* (t_11_ = −3.06, *p* = .011; Figure 2B). The same effect was also present for the alternative measure of metacognitive efficiency *M_diff_*(*= meta-d’ - d’;* t_11_ = −2.99, *p* = .012). Note that while this effect was predicted by our hierarchical model (Figure 1B), it cannot be accounted for by a standard model with no metacognitive noise (Figure 1C).

Further, we examined whether the *M_ratio_* decrease was indeed due to the decrease in sensory noise or to some nonspecific effect of training. We found that subjects who showed a larger decrease in *M_ratio_* also exhibited a larger decrease in intensity threshold (r = .62, *p* = .03; Figure 2C) and a larger increase in *d’* values (r = −.74, *p* = .005; Figure 2D), thus indicating that the *M_ratio_* decrease is directly related to the change in performance on the perceptual task.

Further, one may worry that *M_ratio_* has an intrinsic negative relationship with stimulus sensitivity *d’* independent of sensory noise. To check for this possibility, we computed *d’* and *M_ratio_* across all seven sessions for the lower vs. upper half of intensities used. We found that higher intensities led to a significantly higher *d’* (average *d’* = 2.85 and 0.82 for the upper and lower intensity halves, respectively; t_11_ = 46.23, *p* = 5.9*10^−14^) but did not affect *M_ratio_* (average *M_ratio_* = .98 vs. 1.02 for the upper and lower intensity halves, respectively; t_11_ = −.38, *p* = .71; Figure 2E). Thus, the training-induced decrease in *M_ratio_* cannot be explained as trivially arising from the corresponding *d’* increase.

### Experiment 2: Brief learning leads to lower across-subject metacognitive efficiency

Experiment 1 provided strong support for a causal link between decreased sensory noise and decreased metacognitive efficiency. It employed a standard perceptual learning design with extensive training over a number of days. In Experiment 2 we tested whether much shorter learning period can also lead to decreased metacognitive efficiency. To this end, we recruited a large number of subjects (N = 178) to complete 97 trials of two different perceptual tasks. Critically, we inverted our analyses: rather than combining many trials for each subject (the standard way of analyzing psychophysics data), we combined the data across subjects for a given trial (Figure 3A). This approach allowed us to track the evolution of across-subject performance in terms of both stimulus sensitivity and metacognitive efficiency. Subjects engaged in coarse discrimination of low-contrast Gabor patches (Figure 3B) and fine discrimination on high-contrast Gabor patches (Figure 3C).

**Figure 3.**
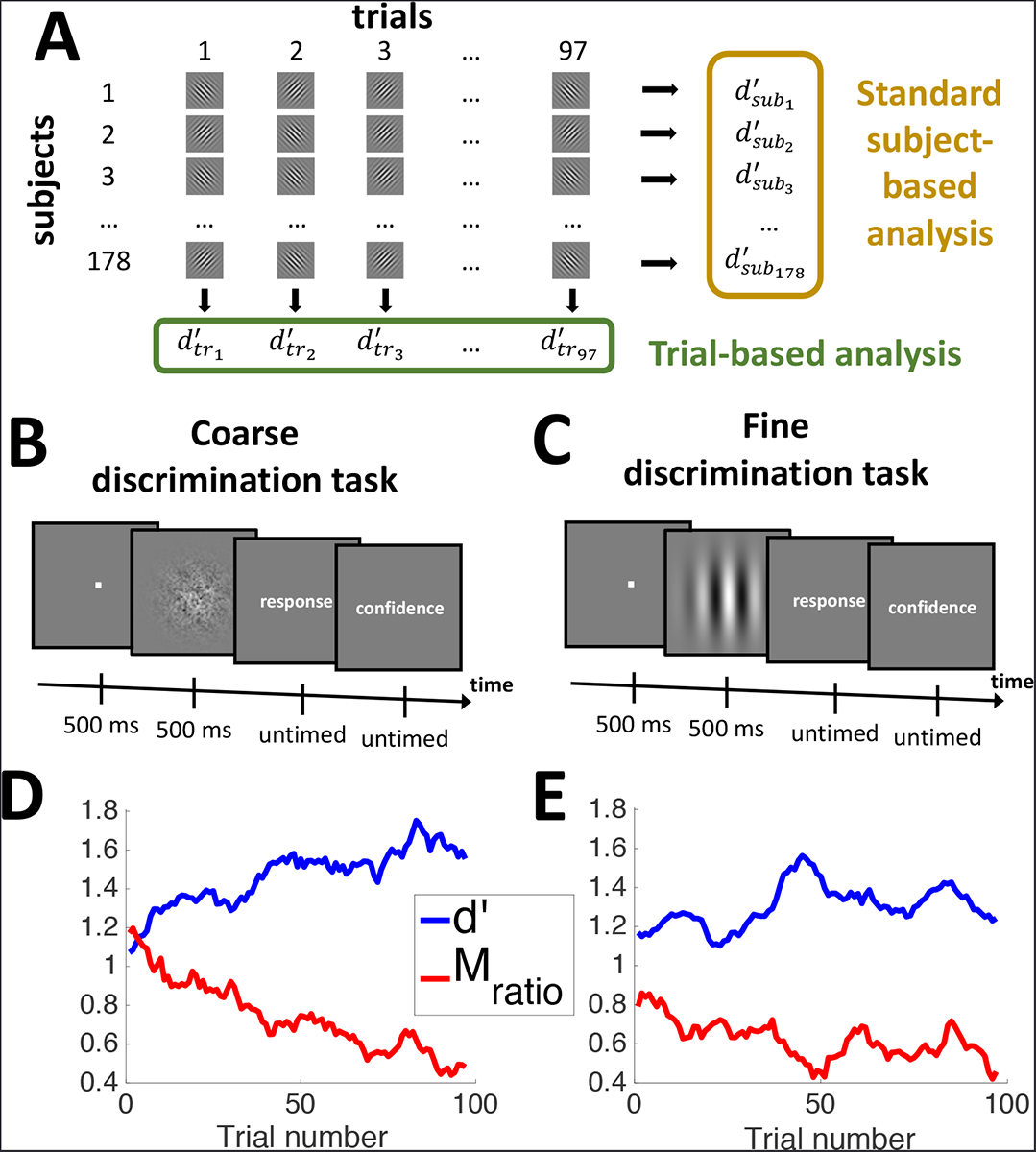
Visual training decreases across-subject metacognitive efficiency. A. Depiction of standard subject-based analysis techniques (which depend on considering all data for a given subject) and trial-based analysis (which depends on considering all data for a given trial number). We investigated the evolution of the trial-based d’ and M_ratio_- B-C. Depictions of the two tasks. Subjects indicated the tilt (clockwise or counterclockwise from vertical) of a Gabor patch and provided a confidence rating on a 4-point scale. In the coarse discrimination task (B), the stimulus was a Gabor patch of low contrast but large tilt (+/-45°). In the fine discrimination task (C), the stimulus was a Gabor patch of high contrast but small tilt. D-E. Practice resulted in a gradual increase in stimulus sensitivity d’ but a decrease in M_ratio_. Both of these effects were larger for the coarse (D) compared to the fine (E) discrimination task. The timecourses are smoothed with a 11-point moving window for display purposes.

As expected, stimulus sensitivity *d’* increased over the course of the 97 trials for both of our tasks (coarse discrimination task: t_95_ = 5.26, *p* = 8.8*10^7^; fine discrimination task: t_95_ = 2.34, *p* = .02; t-tests on the slope parameter in a linear regression; Figure 3D-E). Critically, as in Experiment 1, we observed a corresponding decrease in *M_ratio_* (coarse discrimination task: t_95_ = −6.28, *p* = 9.9*10-9; fine discrimination task: t_95_ = −2.31, *p* = .02; t-tests on the slope parameter in a linear regression; Figure 3D-E).

As can be seen in Figures 3D-E, the learning rate was different for the two tasks. Indeed, the *d’* increase was steeper for the coarse discrimination than for the fine discrimination task (t_190_ = 2.53, *p* = .01). Importantly, we observed a corresponding effect in *M_ratio_*, which showed a steeper decrease for the coarse than the fine discrimination task (t_190_ = −2.85, *p* = .005), suggesting a direct relationship between the amount of learning and the decrease in metacognitive efficiency. All effects pertaining to *M_ratio_* remained significant with the alternative measure of metacognitive efficiency *M_diff_*.

### Experiment 3: Experimentally increasing sensory noise leads to higher metacognitive efficiency

The results of Experiments 1 and 2 lend strong support for the notion that training-induced decrease in sensory noise leads to a corresponding decrease in metacognitive efficiency. Nevertheless, it remains possible that the results of both experiments depended on the use of training and that other manipulations of sensory noise would not produce equivalent results.

To investigate the influence of sensory noise independent of visual training, in Experiment 3 we manipulated the level of sensory noise directly. To do so, we used three levels of contrast and combined them in different ways to construct four conditions that vary on the amount of trial-to-trial variability in the perceptual signal. Twelve subjects performed a Gabor patch orientation discrimination task (Figure 4A) and completed 4,200 trials over the course of three testing days. The Gabor patches were presented with three different levels of contrast. By combining more and more dissimilar contrasts in the same analysis, we constructed four different levels of increasing across-trial stimulus variability (Figure 4B).

**Figure 4.**
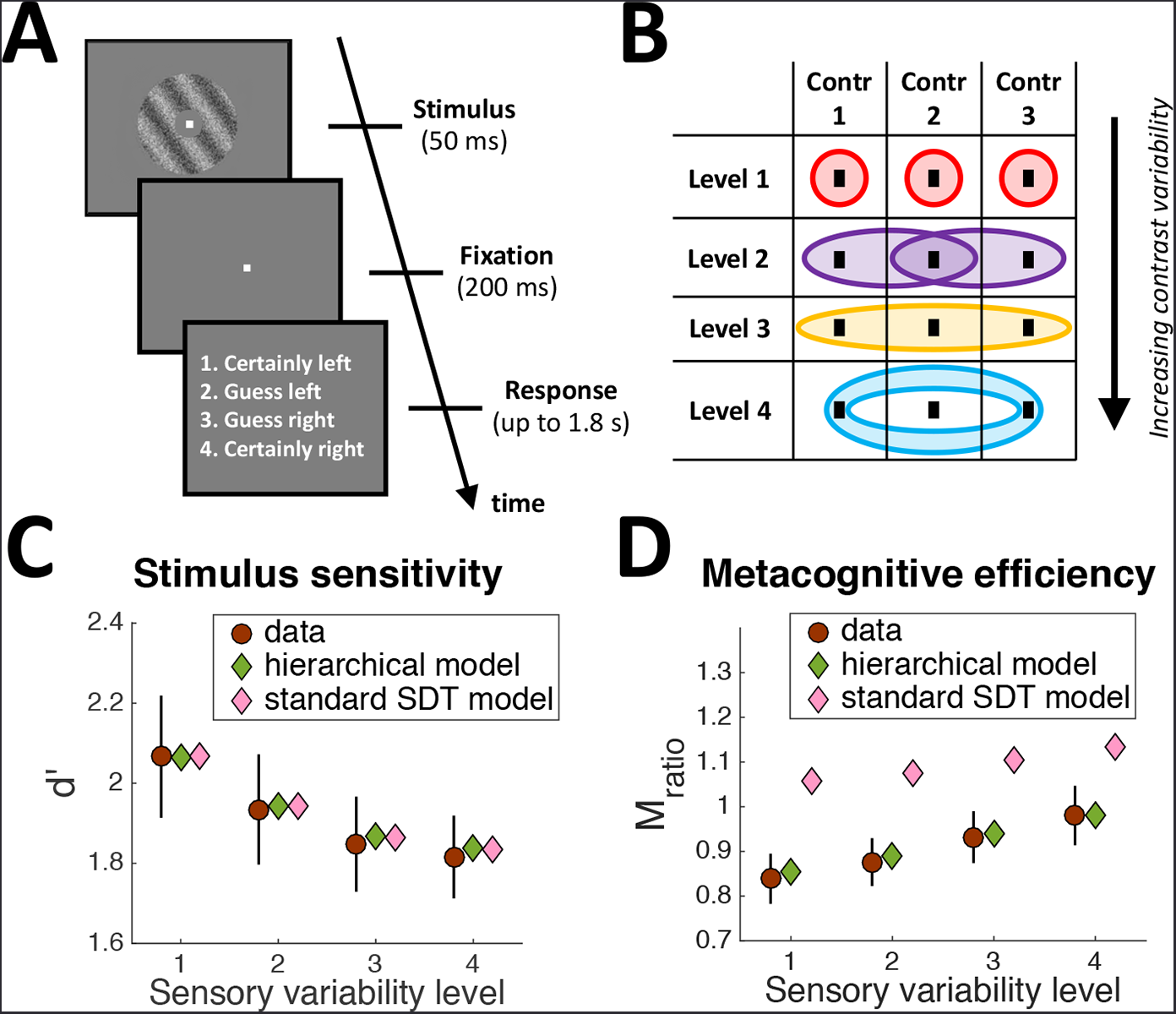
Experimentally increasing sensory noise increases metacognitive efficiency. A. Subjects indicated the tilt (clockwise or counterclockwise from vertical) of a noisy Gabor patch and provided a confidence rating (on a 2-point scale) using a single button press. B. Analysis logic. Three contrast levels were interleaved during the experiment. Different combinations of these contrasts resulted in different levels of stimulus variability. At the lowest level of variability (Level 1), each contrast was analyzed separately and the resulting d’ and M_ratio_ values were averaged. At the next variability levels, increasingly disparate contrasts were combined: nearby contrast pairs in Level 2, all contrast levels in Level 3, and the far-contrasts pair in Level 4. The increased variability in stimulus contrast induced increased sensory variability. C. The four levels of contrast variability were associated with a decreasing stimulus sensitivity d’. This effect was well captured by both our hierarchical model and a standard SDT-based model. D. Higher stimulus variability led to higher metacognitive efficiency M_ratio_. This effect was captured by our hierarchical model but not by the standard SDT-based model. Note that the SDT model predicts both higher M_ratio_ values and a shallower slope of M_ratio_ increase. Error bars indicate S.E.M.

We found that higher levels of stimulus variability led to a decreased *d’* (t_11_ = 4.53, *p* = .0009; Figure 4C). This result may appear surprising since the different conditions consisted largely of the same actual trials that were simply combined in different ways. The robust but relatively modest decrease in *d’* can be explained by the nonlinear relationship between accuracy and *d’* (a detailed explanation can be found in Supplementary Figure 1). Indeed, both our hierarchical and a SDT-based model (see Figure 1C) could capture this decrease (Figure 4C).

Critically, higher levels of across-trial stimulus variability led to an increased *M_ratio_* (t_11_ = 6.21, *p* = .00007; Figure 4D; same effect was observed for *M_diff_ too*, t_11_ = 5.85, *p* = .0001). This effect was quantitatively accounted for by our hierarchical model but not by the standard SDT model (Figure 4D). Most saliently, the SDT model predicted overall higher *M_ratio_* values (average difference = 0.22, t_11_ = 6.06, *p* = .00008). Note that even without metacognitive noise, the SDT model predicts increasing *M_ratio_* values for higher levels of stimulus variability. The reason is that combining disparate contrast values results in violations of the Gaussian variability assumption and this violation is greater for the higher variability levels. Nevertheless, the increase of *M_ratio_* that can be attributed to violations of the Gaussian assumption is smaller than the increase in the data. Indeed, the SDT model predicted a shallower slope of increasing *M_ratio_* values (.026 in model vs. .048 in data, t_11_ = 4.84, *p* = .0005), indicating that metacognitive noise is needed to explain both the lower *M_ratio_* values and the steep *M_ratio_* increase caused by increased stimulus variability.

Since the hierarchical model was more complex than the SDT one (it had one more free parameter), we compared the Akaike Information Criterion (AIC) for each models fit. AIC measures the quality of the fit while punishing for the number of parameters. The hierarchical model still significantly outperformed the SDT model (average AIC difference across the 12 subjects = 23.48 signifying that the hierarchical model is 1.3*105 more likely than the SDT model).

Importantly, as in Experiment 1, we confirmed that simply increasing *d’* does not necessarily lead to a decrease in *M_ratio_*. To demonstrate this point, we analyzed each level of contrast separately and found that higher contrast levels led to higher *d^’^* (*d′_contrastl_* = 1.06, *d′_contrast2_* = 1.9, *d′_contrast1_* = 3.21; slope was significantly positive, t_11_ = 12.9, *p* = 5.5*10^−8^) but without decreasing significantly *M_ratio_* (*M_ratio__contrast1_* = .87, *M_ratio__contrast2_* = .84, *M_ratio__contrast3_* = .81; slope was not different from 0, t_11_ = −1.04, *p* = .32). Further, the *d’* increase from the lowest to highest contrast (Δ *d’* = 2.16) was much higher than between the lowest and highest variability level in Figure 4C (Δ *d’* = .25; t_11_ = 16.49, *p* = 4.2*10^−9^), indicating that the effects in Figure 4D cannot be simply due to the difference in *d’*.

Having confirmed that increasing external stimulus variability in Experiment 3 resulted in increased *M_ratio_*, we looked for a similar effect in Experiment 1. We took advantage of the fact that Experiment 1 included a range of intensity levels and examined the effect of selecting increasingly larger ranges of intensity values. We created four ranges (35-65, 25-75,15-85, and 5-95 percentile of all intensities used) and found that larger ranges did not change *d’* (t_11_ = 1.53, *p* = .15) but led to significantly higher *M_ratio_* values (t_11_ = 5.004, *p* = .0004; Supplementary Figure 2), thus mirroring the effects from Experiment 3.

## Discussion

We found that increasing the levels of sensory noise increases metacognitive efficiency. This effect was robust across experiments and manipulations. The increase of metacognitive efficiency with higher sensory noise was predicted by our hierarchical model of confidence generation that posits a stepwise organization of information flow for perceptual decisions and confidence. Conversely, a standard model based on signal detection theory and lacking independent metacognitive noise could not explain our results. These findings demonstrate the possibility of directly manipulating subjects metacognitive efficiency and provide strong evidence for the notion that confidence ratings are based on different information than perceptual decisions.

A hierarchical model of confidence generation motivated our studies and provided excellent fit to the data. The model assumes that the information available for metacognition is corrupted by extra noise compared to the information available for the perceptual decision. Several previous papers have proposed similar architecture (Berg & Ma, 2016; De Martino et al., 2013; Jang et al., 2012; Maniscalco & Lau, 2016; Mueller & Weidemann, 2008; Rahnev et al., 2016). Here we tested a strong, and previously unrecognized, prediction of hierarchical models on the relationship between sensory noise and metacognitive efficiency. While previous work included metacognitive noise purely to improve model fit, we tested a direct prediction of hierarchical models. Therefore, our results provide some of the strongest evidence to date for the existence of independent metacognitive noise.

An important question concerns how this metacognitive noise can be manipulated directly. Previous research has demonstrated that metacognitive efficiency is affected by fatigue (Maniscalco, McCurdy, Odegaard, & Lau, 2017), working memory demands (Maniscalco & Lau, 2015), and can be enhanced pharmacologically via noradrenaline blockade (Hauser et al., 2017). All of these previous findings rely on taxing subjects “resources” for metacognition. Our findings demonstrate that hierarchical models of confidence can also be used to predict how metacognitive efficiency depends on low-level stimulus characteristics independent of high-level resources.

We modeled the effects of visual perceptual learning as a simple decrease in sensory noise. There is indeed ample evidence that perceptual learning leads to noise attenuation (B. A. Dosher & Lu, 1998, 1999; B. Dosher & Lu, 2017; Petrov, Dosher, & Lu, 2005; Raiguel, Vogels, Mysore, & Orban, 2006). However, at the same time, perceptual learning may also increase the signal (Solovey, Shalom, Pérez-Schuster, & Sigman, 2016), sharpen the perceptual template used to process the stimulus (Li, Levi, & Klein, 2004), improve probabilistic inference (Bejjanki, Beck, Lu, & Pouget, 2011), etc. (for reviews, see Dosher and Lu, 2017; Lu et al., 2011; Watanabe and Sasaki, 2015). Perceptual learning likely has many consequences and our experiments were not designed to distinguish or weight the importance of each of the above effects. Rather, perceptual learning was used as a tool that allowed us to decrease sensory noise in our model. Several previous studies have combined confidence ratings and perceptual learning (Guggenmos, Wilbertz, Hebart, & Sterzer, 2016; Schwiedrzik, Singer, & Melloni, 2011; Solovey et al., 2016; Zizlsperger, Kümmel, & Haarmeier, 2016) but while they found important effects of learning on the overall confidence level, none investigated how training affects metacognitive efficiency.

Our finding of a positive relationship between sensory noise and metacognitive efficiency raises the question as to how metacognitive scores should be interpreted. Influential theories pose that metacognition stems from second-order monitoring processes (Shimamura, 2000). The contents of these second-order metacognitive processes are often assumed to reflect the contents of consciousness (Kunimoto, Miller, & Pashler, 2001; Persaud et al., 2011). However, our results demonstrate that while metacognitive judgments may indeed be related to consciousness, they cannot generally be used as a direct measure of consciousness (Jachs, Blanco, Grantham-Hill, & Soto, 2015). Indeed, perceptual learning has been argued to increase consciousness (Schwiedrzik et al., 2011) but, as seen here, decreases metacognitive efficiency. We see metacognitive scores as invaluable in constructing and testing models of decision making but remain agnostic about their relationship to constructs such as consciousness and working memory.

An important question for future research is whether metacognitive efficiency can be trained. Given that subjects completed the same metacognitive task for seven days, one may expect that their metacognitive noise would decrease. Our design did not allow us to separate the effects of training on sensory and metacognitive noise but given the decrease of metacognitive efficiency, putative decreases in metacognitive noise must have been small. Importantly, we did not include trail-to-trial feedback; such feedback may be more important for decreasing metacognitive compared to sensory noise.

In conclusion, we showed the existence of a robust positive relationship between the level of sensory noise and metacognitive efficiency. These results point to the existence of independent metacognitive noise and have strong implications about the meaning and interpretation of metacognitive efficiency.

## Methods

### Subjects

A total of 225 subjects participated in the three experiments (12 in Experiment 1, 201 in Experiment 2, and 12 in Experiment 3). Each subject participated in only one experiment. Experiments 1 and 3 were conducted in a traditional laboratory setting, while Experiment 2 was conducted online with subjects recruited using Amazons Mechanical Turk. In Experiment 2, subjects who had performed at chance level or failed to clear our attention checks were excluded from the analyses (see below for details). All procedures were approved by the local Institutional Review Board committee. Subjects reported normal or corrected-to-normal vision and provided informed consent.

### Experiment 1

Subjects performed a 2-interval forced choice (2IFC) orientation detection task. Two stimuli were shown in quick succession and subjects indicated the interval (first or second) that contained the target (Figure 2A). The target was a Gabor patch of a particular orientation (circular diameter = 5°, standard deviation of Gaussian filter = 2.5°, spatial frequency = 1 cycle/degree, random spatial phase). The Gabor patch was superimposed by noise generated from a sinusoidal luminance distribution. We varied stimulus intensity by controlling the ratio of noise pixels. The non-target consisted of the superimposed noise only. The target interval was determined randomly on each trial. The center of the Gabor patch was positioned 4° away from the center of the screen in a direction of 45° toward either lower left or lower right. Each trial started with a 500-ms fixation period. The two stimulus intervals lasted 50 ms each, separated by a 300-ms blank period (Figure 2A). Subjects were asked to make two responses: first, to indicate the target interval, and second, to indicate their confidence level. Once the first response was made, the central fixation dot changed color from white to green to signal that the response has been recorded and to cue the need to make a second response. Subjects indicated their confidence using a 4-point scale.

We trained subjects on a specific visual quadrant (either lower left or lower right) and a specific orientation (either 10° or 70°). The trained quadrant and orientation were determined randomly for each subject. Sessions 1 and 7 included testing on the untrained quadrant and orientation (data not reported here). Subjects completed 12 blocks of trials. Each block involved a 2-down 1-up staircase procedure that continuously adjusted the stimulus intensity and terminated after 10 reversals. The intensity threshold for each block was calculated as the geometric mean of the last six reversals per block. In sessions 2-6, all 12 blocks came from the trained condition, while in sessions 1 and 7, four blocks were presented from each of the trained and two untrained conditions (in a randomized order). To keep the sessions as equivalent as possible, data analyses included all four blocks from the trained condition in sessions 1 and 7, as well as the first four blocks in sessions 2-6. Stimuli were generated using Psychophysics Toolbox (Brainard, 1997) in MATLAB (MathWorks, Natick, MA) and were shown on a LCD display (1024 × 768 pixel resolution, 60 Hz refresh rate).

### Experiment 2

Subjects performed two separate tasks - coarse and fine discrimination - that involved discrimination between clockwise and counterclockwise oriented Gabor patches (circular diameter = 1.91°). In the coarse discrimination task (Figure 3B), the stimulus was a Gabor patch of large tilt (+/-45°) overlaid on a noisy background composed of uniformly distributed intensity values. In the fine discrimination task (Figure 3C), the stimulus was a Gabor patch of small tilt (less than 1°) presented without any additional noise.

Each trial started with a fixation cross appearing at the center of the screen. The first trial of each block had was preceded by a longer fixation period of two seconds to allow the subjects time to focus. All other trials had a variable fixation period that was sampled from a uniform distribution with a range of 300-700 ms. The stimulus was then presented for 500 ms. Once the Gabor patch disappeared, subjects were asked to make two responses using their keyboard: first to indicate the tilt of the stimulus and second to rate their confidence on a 4-point scale.

We collected data from three batches of 50 subjects and one batch of 51 subjects. In order to ensure similar average performance on both tasks, we varied the difficulty of each task across the batches. For the coarse discrimination task, difficulty was manipulated by adjusting the contrast level (mean contrast = 5.25%, SD = 0.7%). For the fine discrimination task, difficulty was manipulated by changing the offset from the vertical (mean = 0.69°, SD = 0.09°). Average accuracy was 76.44% for the coarse discrimination task and 74.12% for the fine discrimination task.

Subjects had to complete a total of 100 trials of each task. Each task was divided into five blocks of 20 trials each. Subjects were allowed to take breaks between each block and the order of the tasks was randomized across subjects.

To ensure high data quality, we included six attention check trials - three in each task. These trials were designed to be much easier than the regular trials (contrast for coarse discrimination task = 15%, offset for the fine discrimination task = 5°) and subjects paying attention to the task were expected have a high degree of accuracy for such trials. Therefore, we excluded subjects who responded incorrectly to more than two out of six catch trials (total 15 excluded). Additionally, we excluded subjects whose performance was close to chance level (< 55% correct) on the non-catch trials of either task (additional 8 subjects excluded). These criteria led to the exclusion of a total of 23 of the initial 201 subjects (11% exclusion rate). Note that the final analyses were based only on the 97 non-catch trials per task.

The Gabor stimuli were generated online via in-house code written in JavaScript and the experiment was designed using the JSPsych 5.0.3 library. To account for variability in the resolution and size of screens across subjects, subjects were asked to adjust the size of images of real life objects displayed on the computer screen to match their dimensions to the actual objects. This calibration ensured that the size of the stimulus displayed was uniform across different screens.

### Experiment 3

This study was originally reported as Experiment 2 in Rahnev et al. (2013). All study details can be found in the original publication. Briefly, subjects task was to indicate the tilt (clockwise or counterclockwise) of a grating presented at fixation. Each trial began with 50 ms presentation of the grating followed by a fixation period of 200 ms (Figure 4A). On each trial, the orientation of the grating was randomly selected to be tilted 10° clockwise or 10° counterclockwise away from vertical. The grating pattern was presented on an annulus (inner circle radius: 1.5°, outer circle radius: 4.5°) region. The stimulus consisted of a noisy background composed of uniformly distributed intensity values on top of which we added a grating (0.5 cycles/degree). Subjects were required to fixate on a small white square for the duration of the experiment. They were seated in a dim room 50 cm away from a computer monitor. Stimuli were generated using Psychophysics Toolbox (Brainard, 1997) in MATLAB (MathWorks, Natick, MA) and were shown on a MacBook (13 inch monitor size, 1200 × 800 pixel resolution, 60 Hz refresh rate).

After each stimulus presentation, subjects used one of four keys to give their response indicating the perceived orientation of the grating and a wager on whether they were correct. Subjects used the keys 1-4 indicating “certainly left”, “guess left”, “guess right”, and “certainly right,” respectively. A correct “certain” (i.e., high confidence) choice was awarded with two points while a correct “guess” (i.e., low confidence) choice was awarded with one point. An incorrect “guess” (i.e., low confidence) choice resulted in no points being won or lost but an incorrect “certain” (i.e., high confidence) choice resulted in a loss of two points. We chose this point structure to ensure that subjects gave a sufficient number of both “guess” and “certain” responses. The optimal strategy for this payoff structure was to choose the “certain” choice only when the probability of being correct exceeded 66.7%. We informed subjects of this contingency in order to guarantee that all subjects were aware of the optimal strategy. To further encourage optimal usage of the wagers, we gave the two subjects with highest final scores an additional cash prize. Since the wagers that subjects used were a proxy for their confidence on each trial, for simplicity we refer to the wagers as confidence ratings in the rest of the manuscript.

Each trial lasted for two seconds. Subjects had 1.8 seconds to give their response after the onset of the stimulus. Once a response was given, the text indicating the four possible answers disappeared and the next trial started. If a response was not given in the 1.8-second period, subjects were penalized by a subtraction of four points and the text was removed at the end of the 1.8-second period in order to avoid any potential interference with the processing of the stimulus in the next trial.

The study consisted of four days: one training and three days of testing. In the initial training session on day 1, subjects practiced with the task over the course of five blocks of 120 trials each. Days 2-4 involved theta burst stimulation (TBS) to three different brain areas (visual cortex, Pz, and sham). TBS had a modest effect on subjects performance (reported in the original publication). Here we combined all sessions regardless of TBS condition in order to increase the power of our analyses, which were orthogonal to the TBS effects. Based on the results of the training session on day 1, we chose a grating contrast for each subject that would produce ~80% correct responses. However, we included two more levels of contrast: 75% and 125% of the above contrast. These three contrast levels were used on days 2-4 without further adjustments even if performance deviated from the 80% correct target for the middle contrast. Contrast level was chosen randomly on each trial and subjects were not explicitly informed about the presence of multiple contrast levels. In each session, subjects completed five blocks of 140 trials each for a total of 4,200 trials. Note that the original publication excluded three of the subjects because they did not see phosphenes. These subjects were included here.

### Analyses

To determine observers performance on the task, we computed the signal detection theory (SDT) measure *d’* (a measure of stimulus sensitivity) by calculating the hit rate (HR) and false alarm rate (FAR):

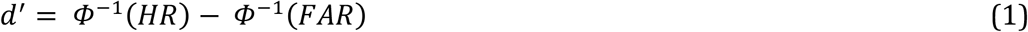

where Ø^−1^ is the inverse of the cumulative standard normal distribution that transforms HR and FAR into *z*-scores. In all experiments, HR and FAR were defined by treating the clockwise orientation as the target. The measures of metacognitive efficiency *M_ratio_* and *M_diff_* were computed using the codes provided by Maniscalco and Lau (2012).

### Model development

Following standard assumptions dating back to the development of signal detection theory (SDT; Green and Swets, 1966), each stimulus category was assumed to produce an internal response corrupted by Gaussian noise. Without loss of generality, we set the counterclockwise stimuli to produce internal response *r_sens_* = 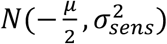 and clockwise stimuli to produce internal response 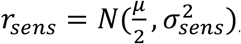, such that the distance between the two distributions was μ. Note that the SDT parameter *d’* can then be expressed as:

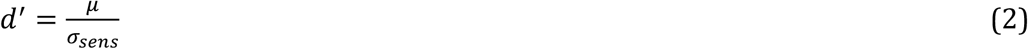

Perceptual decisions were modeled by specifying a decision criterion *c*_0_ and confidence criteria *c*_-n_, *c*_-*n*+1_, …, *c*_−1_, *c*_1_, …, *c_n-1_, c_n_* where *n* = number of confidence ratings. Importantly, the criteria *c*_-n_, *c*_-*n*+1_, … , *c*_−1_ were constrained to be monotonically increasing with *c*_-*n* =_ - ∞ and *c_n =_* ∞. Counterclockwise (clockwise) decisions were made based on whether the internal response *r_sens_* was smaller (larger) than *c*_0_. Confidence responses were given such that an internal response *r_sens_* falling in the internal [*c_i_, c_i+1_*) resulted in a confidence of *i +* 1 when *i* ≥ 0, and of-i when *i* ≤ - 1.

The hierarchical model was constructed similarly but with the important addition of an extra layer of noise. The perceptual decision (about stimulus orientation) was made just as in the standard model described above. However, the confidence judgment was made on the internal signal at a metacognitive stage that was additionally corrupted by Gaussian noise with standard deviation of *σ_meta_* such that signal at the metacognitive stage was given by the formula 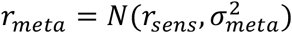. The confidence response was made equivalently to the standard SDT model. However, in cases in which *r_sens_* and *r_meta_* fell on different sides of the decision criterion c_0_, confidence was constrained to always equal 1.

The seven simulations shown in Figure 1B,C were produced by setting σ_*sen*s_ = 1, .83, .7, .6, .55, .52, or .5, while all other parameters were kept constant: *μ* = 1, σ*_meta_* = .3 (in the hierarchical model) or 0 (in the standard SDT model), and the criteria set to *c*_-2_ = — ∞, *c*_2_ = ∞, while *c*_−1_, *c*_0_, and *c*_1_ were set to values corresponding to 30, 50, and 70% posterior probability of a clockwise stimulus. Note that the pattern of results reported in Figures 1B,C is completely insensitive to the exact parameters chosen.

### Prediction of hierarchical models of confidence

Here we give the simple mathematical proof for why hierarchical models of confidence predict that higher sensory noise would lead to higher metacognitive efficiency. As seen in Equation 2, stimulus sensitivity *d’* equals the ratio of the signal and noise present at the decision stage. Equivalently, metacognitive sensitivity *meta-d’* equals the ratio of the signal and noise present at the metacognitive stage. According to our hierarchical model of confidence, the signal at the metacognitive stage is still μ. but the noise is a combination of two Gaussian distributions with standard deviations of σ*_sens_* and σ*_meta_*. Therefore, we can derive that:

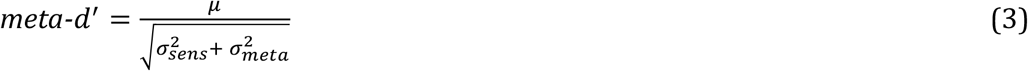

Combining Equations 2 and 3, we obtain:

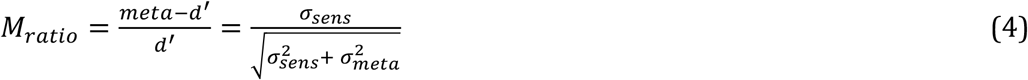

which is an increasing function of *σ_sens_*. Therefore, as sensory noise *σ_sens_* increases, so does metacognitive efficiency *M_ratio_*.

### Model fitting

To model the effect of stimulus contrast in Experiment 3, we set 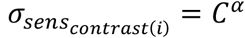, where *C* was set to .75,1, and 1.25 for the three levels of contrast (since contrast levels were 75%, 100%, and 125% of the subject-specific contrast threshold). We do not claim that σ*_sens_* has a power relationship with contrast. However, since this relationship is not generally known, this way of specifying the relationship allowed us to capture any combination of the two ratios between the sensory noise corresponding to successive contrast levels. Importantly, the parameter α was strongly correlated between the fits for the SDT and the hierarchical models (r = .77, *p* = .004), demonstrating that the superior fits of the hierarchical model were not due to an interaction between ***α*** and the extra parameter σ*_meta_*. Finally, since three of the 12 subjects exhibited *M_ratio_* values larger than 1 in at least one condition, we included additional decision-level noise for them and applied it to both the hierarchical and the SDT models.

The SDT and hierarchical models were instantiated with four and five free parameters, respectively. Importantly, the signal μ corresponding to each contrast level was not treated as a free parameter but was directly computed using Equation 2 using the contrast-specific *d’* and sensory noise values. The standard SDT model thus had four free parameters: α and the criteria *c*_−1_*, c_0_*, and *c_1_* (since confidence was provided on a 2-point scale). The hierarchical model was instantiated with five free parameters (the four from the SDT model and σ*_meta_*). The criteria *c_i_* were constrained to be non-decreasing and *σ_meta_* was constrained to be **≥** 0.

We fit the models to the data as previously (Rahnev et al., 2011, 2013; Rahnev, Maniscalco, Luber, Lau, & Lisanby, 2012) using a maximum likelihood estimation approach. The models were fit to the full distribution of probabilities of each response type contingent on each stimulus type. Model fitting was done by finding the maximum-likelihood parameter values using a simulated annealing (Kirkpatrick, Gelatt and Vecchi, 1983). Fitting was conducted separately for each subjects data by first running the fitting five times with general starting parameter set, and then running the fitting five more times using a starting parameter set derived from the best fit from the previous stage. The best fitting model from the second stage was used for further analyses. Akaike Information Criterion (AIC) was used for model comparison though the results remained the same if Bayesian Information Criterion (BIC) was used instead.

## Data and code availability

All data and codes for the analyses are freely available online at https://github.com/DobyRahnev/sensory_noise_metacognitive_efficiency.

## Acknowledgements

We thank Hakwan Lau, Brian Odegaard, and David Soto for helpful comments. This work was funded by a startup grant to D.R. from the Georgia Institute of Technology.

## Supplementary Results

In Experiment 1, we selected a limited range of stimulus intensity values for analyses in order to investigate how training affected stimulus sensitivity, confidence, and metacognitive efficiency. Specifically, we considered all stimulus intensity values used across the seven days and selected all intensities within the 35-65 percentile. We used a relatively small window in order to avoid excessive stimulus variability. However, this choice may appear arbitrary. Therefore, we tested the robustness of our results to using larger ranges of intensity values. In three more analyses, we selected all intensities within 30-70, 20-80, and 10-90 percentile of all intensities. We found that when considering these intensity ranges, we still found the same increase in stimulus sensitivity *d’* (30-70 percentile: t_11_ = 4.92, *p* = .0005; 20-80 percentile: t_11_ = 5.65, *p* = .0001; 10-90 percentile: t_11_ = 7.26, *p* = .00002) and decrease in metacognitive efficiency *M_ratio_* (30-70 percentile: t_11_ = −2.82, *p* = .0165; 20-80 percentile: t_11_ = −2.6, *p* = .0246; 10-90 percentile: t_11_ = −2.66, *p* = .0223). Therefore, our results do not depend on the exact percentile selected.

### Supplementary Figures

**Supplementary Figure 1.**
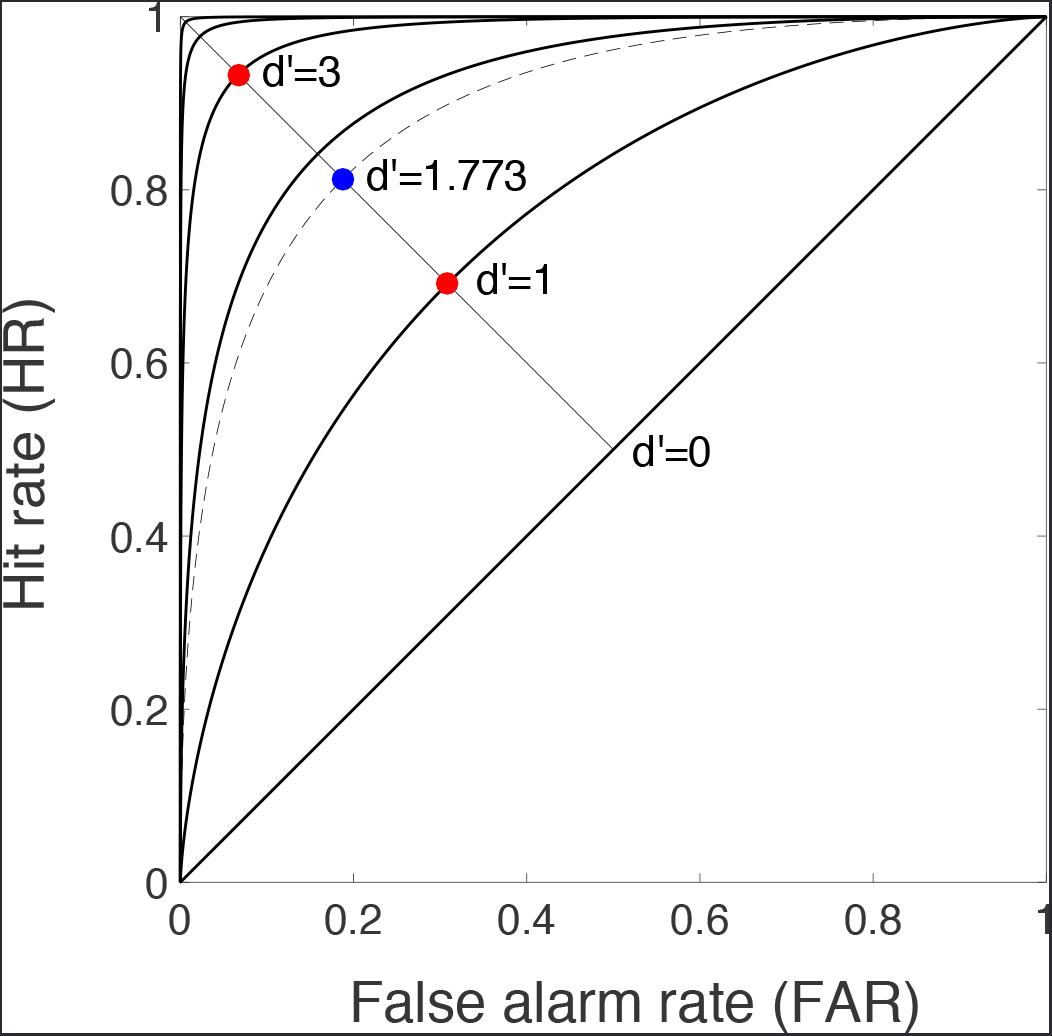
A graphical explanation as to why the *d’* value of two combined conditions is lower than the average *d’* values of those conditions. The figure shows the receiver operating characteristic (ROC) curves for *d’*of 0,1, 2, 3, 4, and 5. The ROC curve plots the false alarm rate (FAR) on the x axis vs. the hit rate (HR) on the y axis. Assume that an unbiased observer (who chooses each stimulus category equally often) shows stimulus sensitivity values of *d’* = 1 and *d’* = 3 in two different conditions. Such performance would result in the red dots marked on the graph, lying on the diagonal perpendicular to the line of *d’* = 0. Assuming that the two conditions had equal number of trials from each category, then their combination would result in a point on the ROC curve (marked in blue) lying exactly midway between the two red circles. As can be seen in the graph, this point corresponds to *d’* of 1.773, which is lower than 2 - the average of 1 and 3. Mathematically, the distance from the *d’* = 0 line (which we can denote with l) is equal to 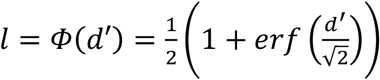, where Φ is the cumulative normal distribution, and *erf* signifies the error function. Therefore, *d’* = Φ^−1^(*l*) and since Φ^−1^(*x*) is convex in [0,1], then 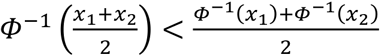 for 0 ≤ *x*_l_ ≤ *x*_0_ ≤ 1, which means that the *d’* of two combined conditions is lower than the average *d’* of each of those conditions. The same arguments hold even if the observer is not unbiased and thus points on the ROC curve do not lie on the diagonal perpendicular to the line of *d’*=0.

**Supplementary Figure 2.**
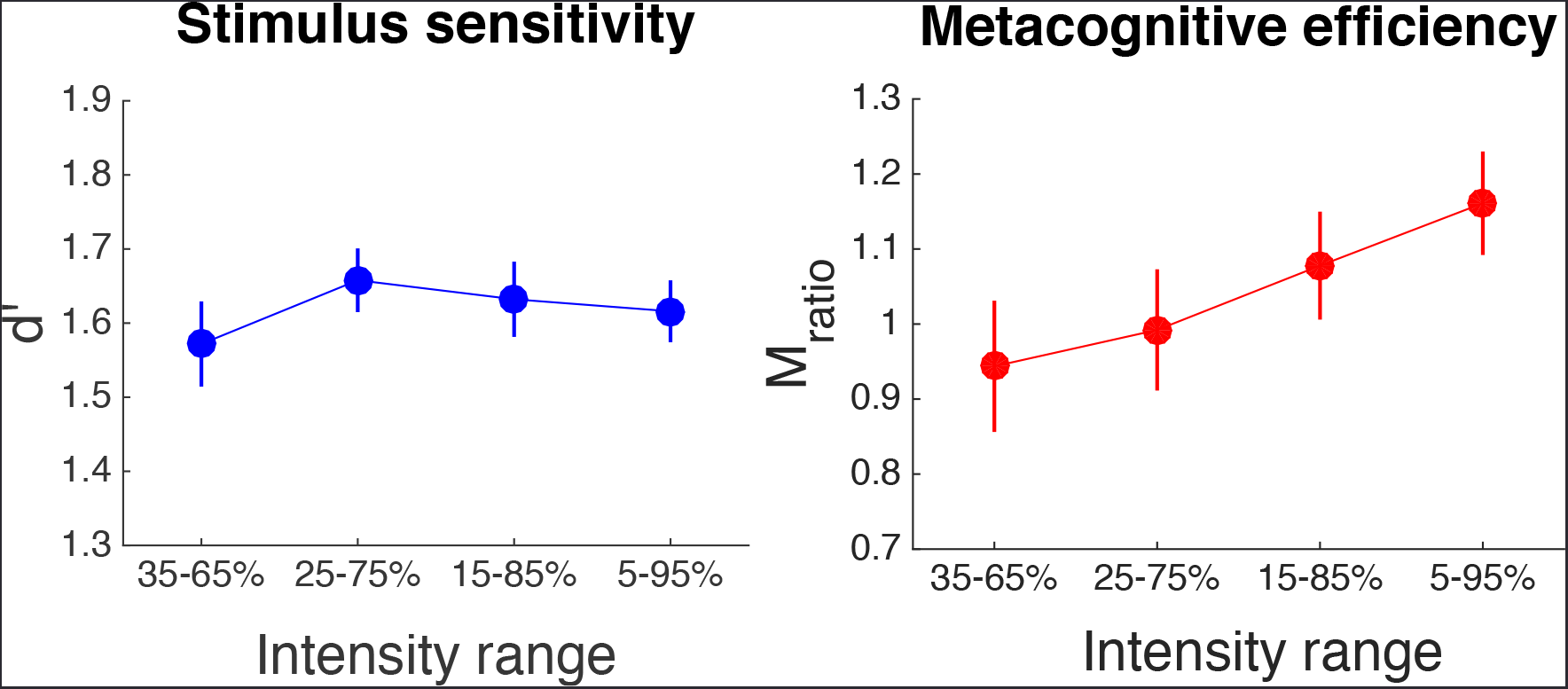
Effects of larger intensity range on *d’*and *M_ratio_*. In Experiment 1, we created four different conditions by choosing intensities that lie within a certain percentile range of all used stimulus intensities. We chose 35-65, 25-75,15-85, and 5-95 percentiles, though different choices result in the same pattern of results. As reported in the main text, larger ranges resulted in slope of *d’* change over the four conditions that is not significantly different from 0 (t_11_ = 1.53, *p* = .15) but led to significantly positive slopes of *M_ratio_* values (t_11_ = 5.004, *p* = .0004), indicating that larger ranges resulted in higher metacognitive efficiency. Error bars represent S.E.M.

